# Quasi-periodic migration of single cells on short microlanes

**DOI:** 10.1101/809939

**Authors:** Fang Zhou, Sophia A. Schaffer, Christoph Schreiber, Felix J. Segerer, Andriy Goychuk, Erwin Frey, Joachim O. Rädler

**Affiliations:** Faculty of Physics and Center for NanoScience, Ludwig-Maximilians-Universität München, Munich, Germany; Arnold-Sommerfeld-Center for Theoretical Physics, Faculty of Physics and Center for NanoScience, Ludwig-Maximilians-Universität München, Munich, Germany

**Keywords:** Cell migration, quasi-periodic, polarization, actin dynamic, micropattern

## Abstract

Cell migration on microlanes represents a suitable and simple platform for the exploration of the molecular mechanisms underlying cell cytoskeleton dynamics. Here, we report on the quasi-periodic movement of cells confined in stripe-shaped microlanes. We observe persistent polarized cell shapes and directed pole-to-pole motion within the microlanes. Cells depolarize at one end of a given microlane, followed by delayed repolarization towards the opposite end. We analyze cell motility via the spatial velocity distribution, the velocity frequency spectrum and the reversal time as a measure for depolarization and spontaneous repolarization of cells at the microlane ends. The frequent encounters of a boundary in the stripe geometry provides a robust framework for quantitative investigations of the cytoskeleton protrusion and repolarization dynamics. In a first advance to rigorously test physical models of cell migration, we find that the statistics of the cell migration is recapitulated by a Cellular Potts model with a minimal description of cytoskeleton dynamics. Using LifeAct-GFP transfected cells and microlanes with differently shaped ends, we show that the local deformation of the leading cell edge in response to the tip geometry can locally either amplify or quench actin polymerization, while leaving the average reversal times unaffected.

## Introduction

Cells navigate in complex environments and undergo morphological changes via dynamic reorganization of the actin cytoskeleton [1, 2]. Movement is generated by cyclic phases of protrusion, adhesion to the extracellular environment, and actomyosin-driven retraction of the cell rear. Actin polymerization and crosslinking prevails in the advancement of filaments, protrusions and lamellipodia. Unraveling the mechanisms underlying actin transport, polymerization dynamics, and their regulation by Rho family GTPases are central challenges towards an intricate understanding of cell migration. The dynamics of actin indeed show many peculiarities, including traveling wave patterns [3-6], retrograde actin flow at the leading edge [2, 7-9], protrusion-retraction cycles as well as persistent polarity [5, 10]. In 2D cell culture, the actomyosin-driven shape changes of the cell body lead to phenotypic migratory modes that can be detected across large length scales. The macroscopically apparent persistent random walk is generated by the following key components: (i) persistence of leading protrusions and (ii) spontaneous front-rear polarization of cells. The cell cytoskeleton that is responsible for cell locomotion is in turn regulated by intracellular signaling proteins like the Rho family of GTPases [11], whose biochemical interactions have been studied both in conceptual and in detailed models [12-18]. In general, the mass-conserving reaction-diffusion systems formed by intracellular proteins can exhibit a wide variety of spatiotemporal patterns [19]. From a theoretical perspective, the formation of such patterns can be understood in terms of shifting local equilibria due to lateral mass redistribution between diffusively coupled reactive compartments [20, 21]. Detailed spatiotemporal models that account for cell shape changes, in response to the formation of Rho GTPase patterns and their regulation of the cytoskeleton, were found to reproduce front-rear polarization of cells [15, 22, 23]. The biophysical principles that underlie the coupling between polarization and migration of cells and determine their shape have been explored by a variety of successful conceptual approaches [24-30]. To rigorously test these models, it is necessary to employ experimental techniques that are capable of studying the shape, migration and internal chemistry of cells in a well-controlled and high-throughput fashion.

In recent years, artificial micropatterns have been used to confine cell migration to well-defined geometries [31], in particular microlanes [32-35]. On microlanes, cell motion is reduced to an effective 1D persistent random walk. There, a universal relation between persistence and cell velocity was shown to hold [7]. Other micropatterns with non-trivial geometries give rise to novel migratory behavior: circular adhesion islands lead to rotational migration of small cohorts of cells [36], ratchet geometries induce directed migration [37-39], cells confined in dumbbell-shaped micropatterns undergo repeated stochastic transitions characterized by intricate nonlinear migratory dynamics [40], and microlanes with gaps show emergence of stochastic cell reversal and transits [41]. In addition, migration patterns may change upon interference with the cytoskeleton. For example, persistent cell migration on linear microlanes shifts to striking oscillations upon depolymerization of microtubules [42] or by depletion of zyxin, a protein that concentrates at focal adhesions and actin cytoskeleton components [43]. Because of their flexibility in controlling cell behavior, micropatterns are well suited to verify computational models of cytoskeleton dynamics and to advance our understanding of the underlying regulatory network. In particular, computer simulations have predicted periodic migration of cells on 1D micropatterns [27]. Similar findings were reported in reaction-diffusion models of actin waves on flexible and on circular boundaries [3, 24]. These theoretical studies suggest that confining geometries might reinforce sustained oscillations. However, systematic experimental studies of periodic cell migration on micropatterns have not yet been carried out. In particular, there have been no studies regarding the dynamics and curvature-dependence of repolarization in the presence of a boundary.

Here, we study the migration of single cells on short microlanes. Using micro contact printing, we create arrays of fibronectin-coated stripe-shaped micropatterns of different lengths. On these micropatterns, we observe quasi-periodic cell migration for the breast cancer cell line MDA-MB-231. We investigate how the spatial distribution of the cell position, the velocity distribution, and the periodicity of cell migration depend on the microlane length. Our data indicate that each cell undergoes repeated cycles of directed migration with pronounced cell polarization, followed by distinct termination of the cell’s leading edge at the micropattern ends, and spontaneous cell repolarization in the opposite direction. We recapitulate these migratory features in a dynamic Cellular Potts model, which includes a simplified description of the adapting cell cytoskeleton [30]. Subsequently, we compare the distributions of apparent repolarization times between our experiments and computer simulations. Finally, we discuss how microlanes with curved ends constitute a controlled experimental framework that enables further investigation into the natureof excitable dynamics in actin-based cell migration.

## Results and discussions

### 1. Single cell migration on stripe-shaped microlanes

In a first set of experiments, we investigated whether cells captured on microlanes exhibit oscillatory migration. Breast cancer cells (MDA-MB-231) are seeded on arrays of fibronectin-coated microlanes, which are surrounded by a PEGylated and therefore cell-repellent surface. These microlanes feature five different lengths between 70 and 270 μm. The fabrication of the micropatterns follows previous protocols and is described in detail in the methods section. Cells adhere, spread and remain confined within the microstructures during the entire period of the experiment. Movies were taken in time-lapse mode, recording images every 10 min over a period of 36 h. During this time, cells migrate in a guided manner and always align their front-rear polarity axis along the main axis of the microlanes. We observe repeated cycles of directed motion, termination of the motion at the micropattern ends, and cell repolarization in the opposite direction. This recurring sequence of events leads to quasi-periodic migration as shown in the time sequences in Fig 1a. Cells exhibit a typical migratory morphology with an actin-rich lamellipodium at the leading edge – seen as a dark rim in phase contrast – and a retracting tail at the rear. As cells reach the respective end of a microlane, they adopt a shortened, almost round appearance with no lamellipodium, until a newly formed lamellipodium appears at the opposite end of the cell body. These migratory and resting phenotypes coincide with distinct regimes of cell motion, which we obtain by tracking the cell nucleus. Fig 1b shows an exemplary trajectory of a cell nucleus over the course of 36 hours. For a video of this exemplary cell, please refer to Video S1 in the Supporting Information. The cell shows phases of directed motion followed by pausing and repolarization, thereby resulting in a quasi-periodic movement. Note that cell motion is quasi-periodic in the sense that the time needed for reorientation of the cell is stochastic, which leads to variability in the period of the back-and-forth motion.

**Fig 1.**
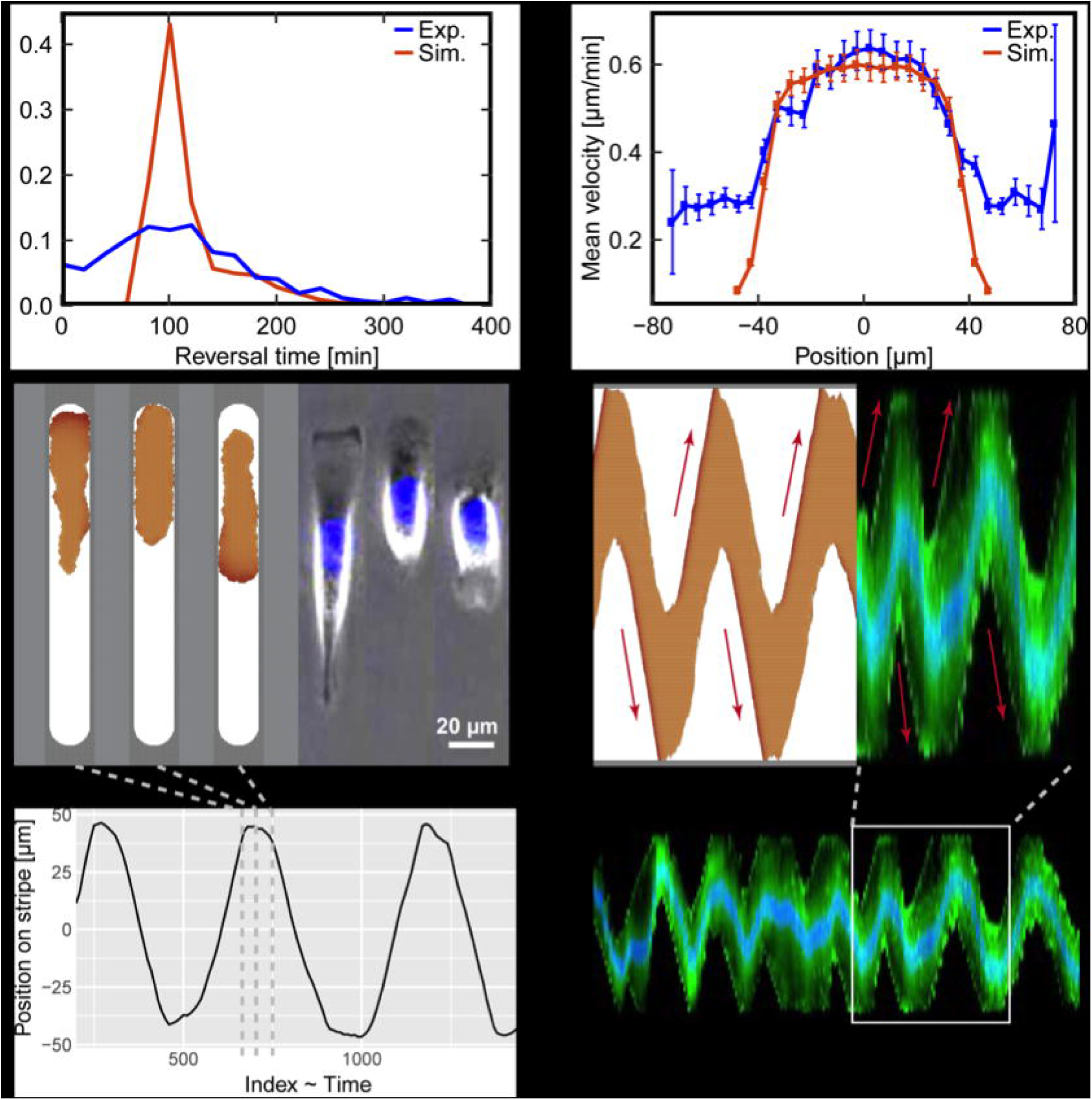
Cell migration on microlanes: (a) Time sequence of a cell (MDA-MB-231) migrating on a stripe-shaped micropattern (Stripe length L = 120 µm, width W = 20 µm, 10 min time intervals). Images are taken in phase contrast and with fluorescence microscopy. The nucleus is labeled with Hoechst 3334 as indicated in blue. (b) Trajectory of the cell nucleus tracked over the course of 36 h showing quasi-periodic alternations between directed migration and repolarization. Additional exemplary trajectories of cells, which represent the broad distribution of frequencies, can be found in Fig. S2 in the Supporting Information.

### 2. Spatial velocity distribution

In order to better distinguish phases of directed migration from phases of reorientation, we fabricated microlanes of different lengths. A sufficiently large sample size of hundreds of cells was acquired by parallel automated tracking of fluorescently labeled cell nuclei. Our automated image analysis yields cell trajectories *x*(*t*) as described in the Materials and Methods. The instantaneous cell velocities are determined by *ν*(*t*) = [*x*(*t* + Δ*t*) – *x* (*t*)] /Δ*t*, where Δ*t* = 10 *min* is the time interval between two subsequent frames. Fig 2 shows exemplary single cell trajectories together with the spatial distributions of cell positions and velocities, which were sampled from ensembles of about 100 cells for each microlane length. These distributions are determined by binning the cell positions into 5 μm wide sections along the microlane, and then computing the fraction *p*(*x*) and the mean absolute velocity ⟨|*ν*|⟩(x) of cells found in each bin. Cells on the shortest microlane (L = 70 μm) do not exhibit periodic motion, and instead remain in a symmetric morphology with two lamellipodia extending at the cell tips. Evidently, there is not enough space for directional migration on short microlanes. In contrast, quasi-periodic migration is observed on longer microlanes (L = 120 - 270 μm). Fig 2b shows that the detection frequency of cell nuclei is flat in the middle of the microlanes and decreases towards the micropattern tips. Note that the spatial distribution of cells in the lanes of length 170 μm shows peaks at both ends, which could be explained by a finite repolarization time that leads to a higher frequency of finding cells at the microlane tips. For longer stripes, the noise level of the spatial distribution of cells is increasing as each point is visited less often, which could prevent the observation of clear peaks there. Similarly, the mean absolute velocity distributions show a distinct plateau behavior in the middle of the microlanes and decline towards the micropattern tips in the case of longer microlanes (220 μm and 270 μm) (Fig 2c). We find that the velocity declines within similarly sized microlane tip regions for all microlane tip lengths, and use this observation to define the onset of cell repolarization. For long microlanes, the spatial velocity distributions appear to have a trapezoidal profile, where two velocity ramps at the microlane tips are connected by a plateau in the microlane center. Therefore, we define each transition point between a ramp (repolarization) and a plateau (run) region as the boundary of the corresponding repolarization region (see black dashed lines in Fig 2). In order to identify these transition points, we apply a change-point-analysis as described in the Supporting Information S2. We find that the distance from the transition points to the tip of the microlane ξ_0_ = 55 μ*m* is nearly constant across all micropattern lengths. Thus, cells on 120 μm stripes are able to polarize but have only a very short migration phase until interacting with the opposite boundary. In the center of longer microlanes (L =170, 220, and 270 μm), we find longer phases of directed motion in which cells migrate with a mean absolute velocity of approximately 0.4 - 0.6 μm/min. Note that the velocity in the middle of the 170 μm microlanes is less uniformly distributed and slightly larger than for the 220 and 270 μm microlanes. However, because cell velocities show large variations, these fluctuations could be caused by the inherent variability of the cells.

**Fig 2.**
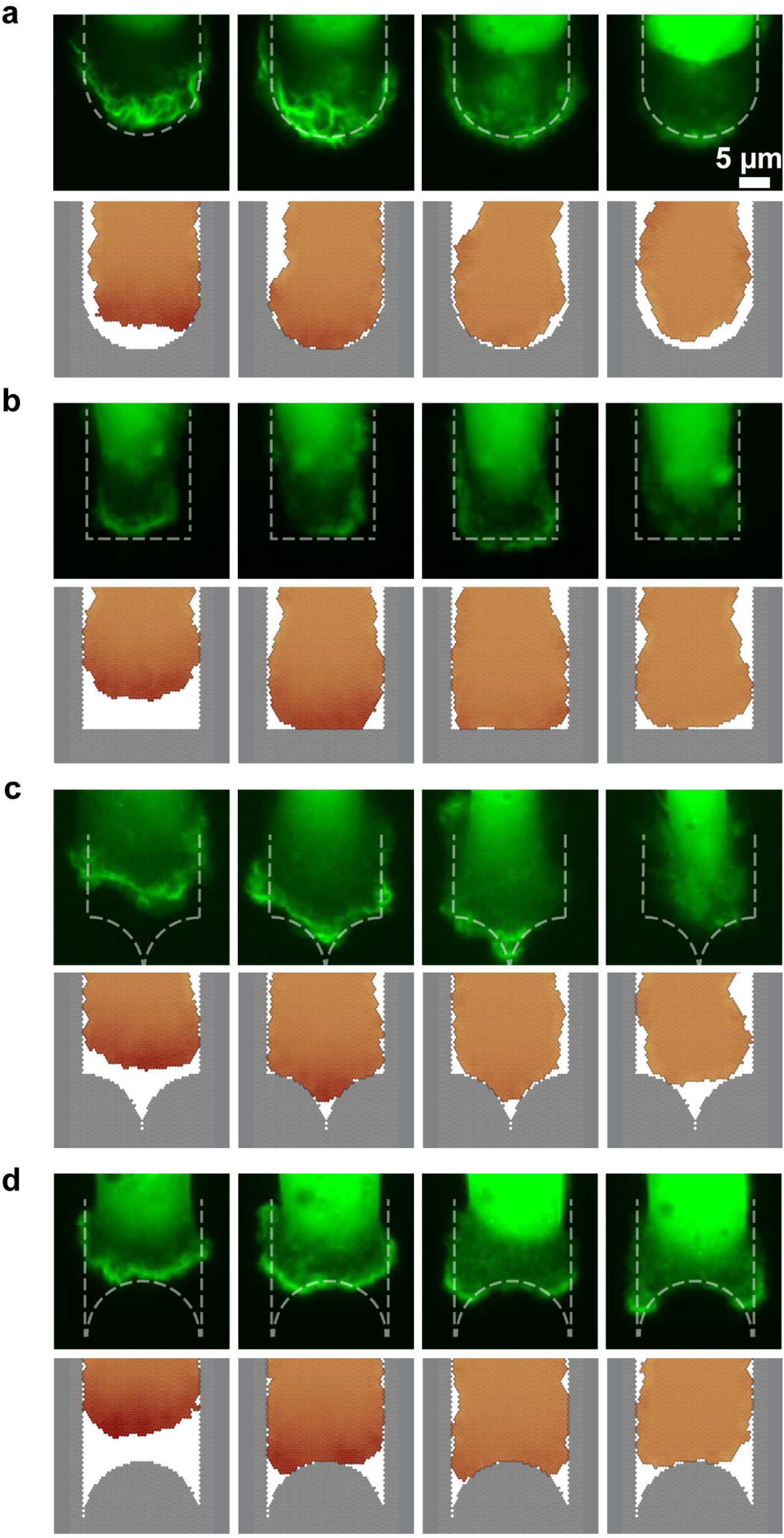
Migration pattern as a function of lane length: From left to right: (a) Time courses of cell nucleus position of cells within microlanes. (b) Spatial distributions of nuclei. (c) Spatial distributions of mean absolute cell velocities. Blue lines represent the standard error of the mean for the binned data. (d) Schematic drawing of the microlanes with length L = 70, 120, 170, 220, 270 μm and width W = 20 μm. Dashed lines indicate the boundary between microlane tips and microlane center. These results were obtained by binning the cell positions (5 μm bin width). For each microlane length, we tracked roughly 100 cells. For videos of exemplary cells on a short microlane (L = 70 μm) and on a longer microlane (L = 170 μm), please refer to Videos S1 and S2 in the Supporting Information.

### 3. Velocity distribution and sustained oscillations

We further quantify the quasi-periodic migration of cells. First, we determine the overall distribution of absolute cell velocities (Fig 3a). We find that for microlanes long enough to show persistent cell migration (L = 120 - 270 μm), the cell velocity distributions collapse onto an exponentially decaying master curve. Short microlanes, where we do not observe persistent cell migration, show a distinctly narrower velocity distribution. This is likely explained by the observation that cell repolarization begins at a distance of 55 μm from the micropattern tips. If the microlane is shorter than twice this distance, then the cell is in a constant state of repolarization, which diminishes oscillatory motion. Subsequently, we performed a discrete Fourier transform of the cell velocity time-traces for different stripe lengths, which yields the frequency distribution corresponding to the quasi-periodic cell migration. We find that the frequency spectrum follows a log-normal distribution. Furthermore, the dominant frequency (*f*_*max*_, peak of the frequency spectrum) shifts to lower frequencies for longer stripe lengths. In the inset of Fig 3b, we find that the period of migration *T* = 1/*f*_*max*_ increases linearly with the stripe length at a slope *dT*/*dL* = 0.054 ± 0.006 h/μm. This indicates that cells move with a constant velocity, *ν*_*c*_ = 0.62 μ*m*/*min*, across the microlane and that their repolarization time does not depend on the length of the microlane. We find that the constant velocity obtained from our spectral analysis is in good agreement with the velocity plateau in the stripe centers (Fig 2). Furthermore, the linear fit intersects the x-axis at *x* = 29.4 μ*m*, which can be interpreted as a lower bound for the microlane length to allow cell motion.

**Fig 3.**
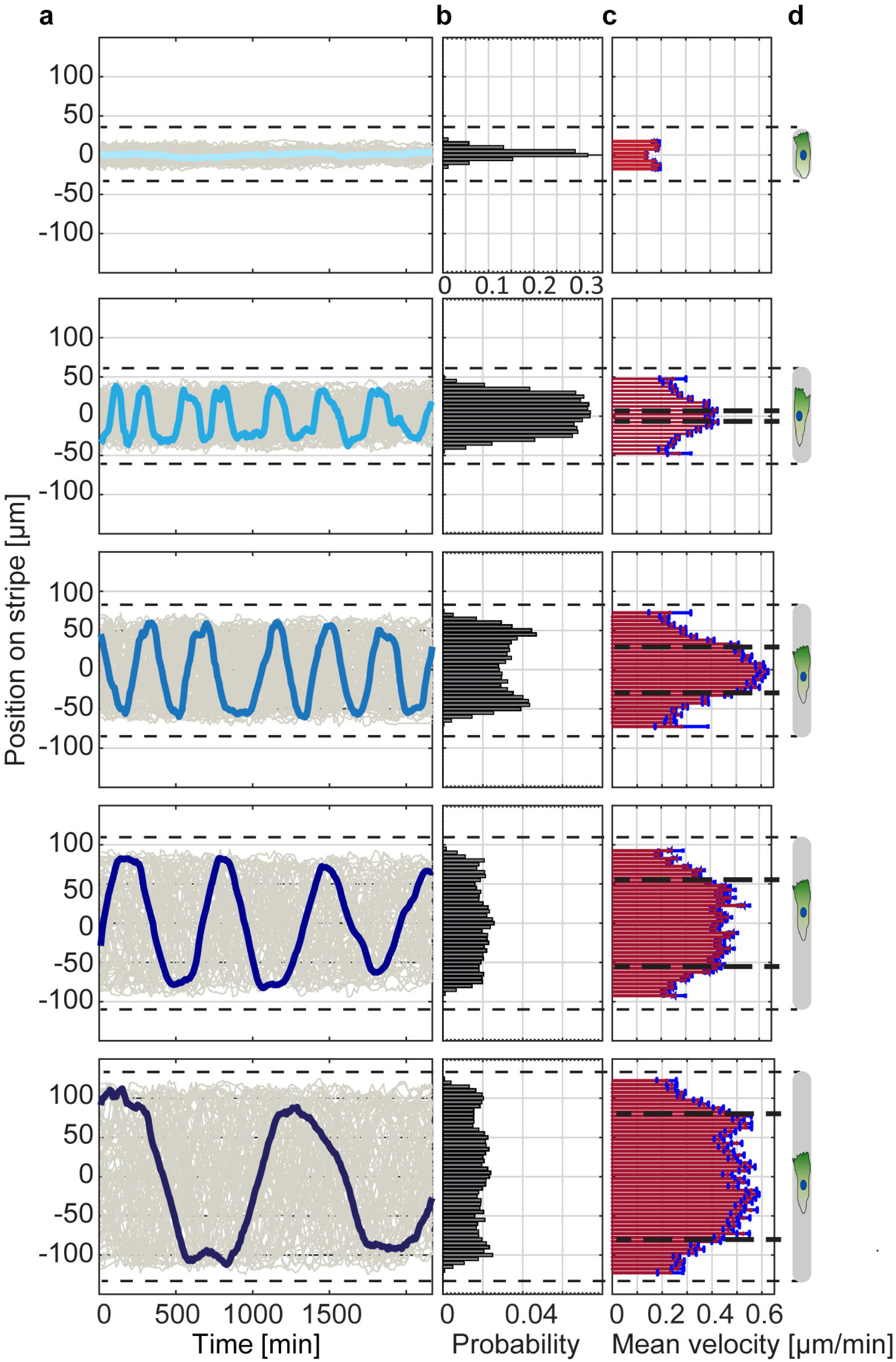
Overall spectral analysis of the large ensemble of cell traces. (a) Normalized distribution of absolute cell velocities, ⟨|*ν*|⟩, for single MDA-MB-231 cells migrating in stripe-shaped micro patterns of five different lengths and fixed width (W = 20 μm). Inset: Non-normalized (counts) velocity distribution in a logarithmic plot. (b) Discrete Fourier transform of the time-dependent directional velocities, averaged over the cell ensemble, fitted by a log-normal distribution (lines). Inset: The migration period (T = 1/frequency) of single cells increases linearly with the stripe length. The error bars correspond to the peak width of the fitting in (b).

### 4. Repolarization time

In the following, we address cell repolarization and the dynamics of directed migration reversal at the ends of the microlanes. We observe that cells depolarize when the protruding lamellipodium encounters the confining PEG-layer. The cell then compacts as its trailing edge continues to move. As the trailing edge stalls, the cell begins to expand again and finally repolarizes towards the opposite, free cell edge. This repolarization manifests itself in a new lamellipodium that emerges at the free cell edge. Note that this phenomenon of internal repolarization is rather specific for cells confined on microlanes. The more general appearance of mesenchymal cell migration in 2D as well as in 3D appears to redirect existing lamellipodia or exhibit several competing lamellipodia. However, in the experiments presented here, the appearance of lamellipodia is restricted to two opposite sides of the cell due to its lateral confinement by the microlanes. Hence, reorientation of crawling cells occurs via a relatively well-defined cycle of depolarization and repolarization. This feature allows us to examine the “reversal time”, which is a measure for the time scale of depolarization and repolarization when a cell reaches the end of a microlane. As discussed in section 2, “Spatial velocity distribution”, we define a reversal area A_0_ at the ends of the microlanes (Fig 4a); in Fig 2 the reversal area marks the regions where we observe a decrease of the cell velocities (with a distance ξ_0_ = 55 μm to the boundary). During a depolarization-repolarization cycle, a cell first enters the reversal area at *t* = *t*_1_ when it approaches the microlane tip, and leaves it at *t* = *t*_2_ after repolarization. Therefore, we define the reversal time as *t*_*R*_ = *t*_2_ – *t*_1_ and determine its distribution for four different microlane lengths (Fig 4b). We note that the reversal time distributions are independent of the stripe length, which is in good agreement with our spectral analysis in section 3, “Velocity distribution and sustained oscillations”. Although the exact value of the average reversal time will typically depend on the particular choice of the reversal area, we consistently find an average depolarization and repolarization time of approximately 100 min.

**Fig 4.**
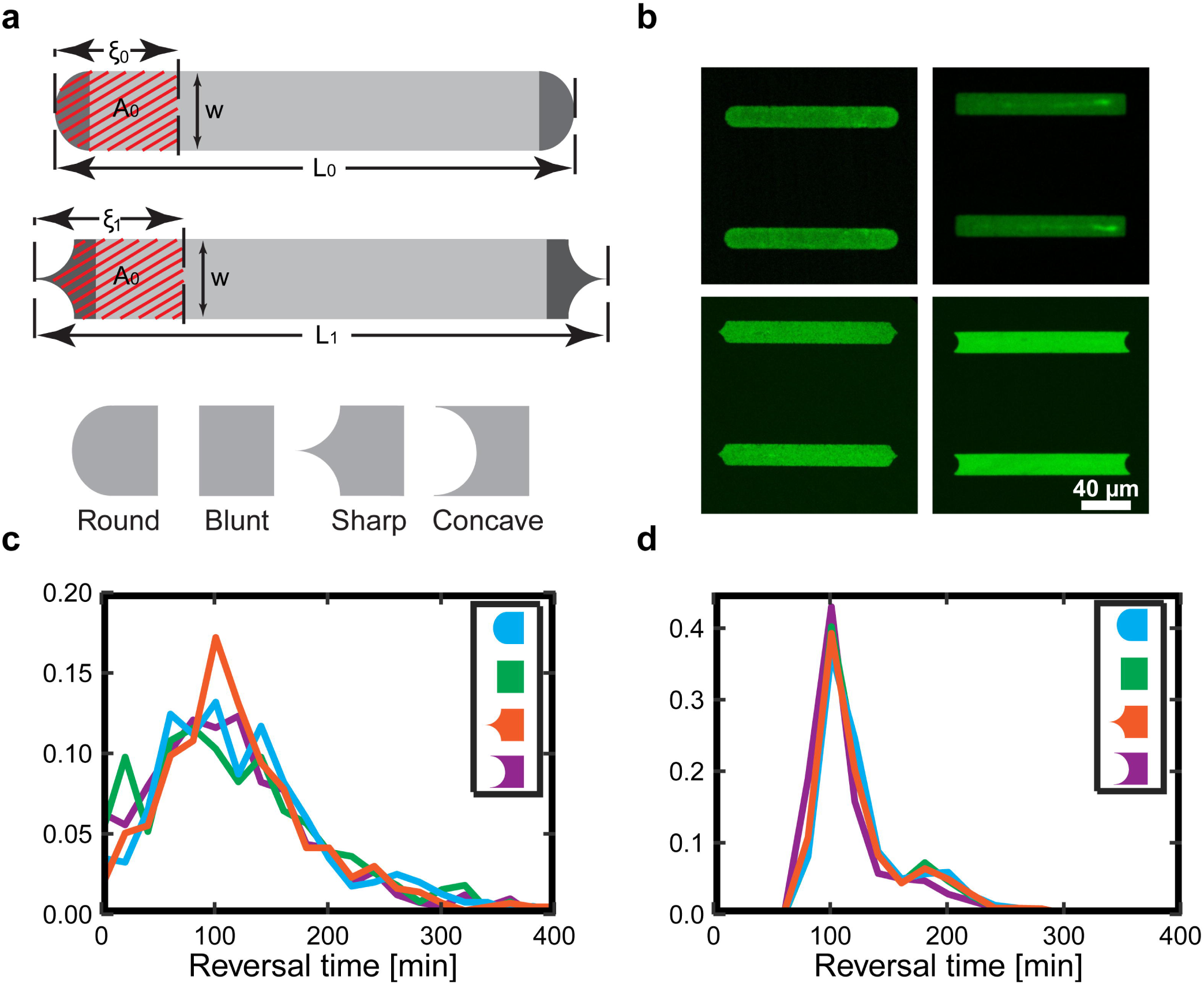
Distribution of “reversal times”: (a) Schematic drawing of a cell entering the microlane tip region A_0_ (red hatches), arresting and then returning from the tip region, with the leading edge of the cell shaded in dark green. The red dashed line indicates the position of cell entry into the tip region A_0_, at time *t*_1_, and exit from the tip region, at time *t*_2_. (b) Normalized distribution of the reversal times *t*_*R*_ = *t*_2_ – *t*_1_ for four different microlane lengths. Inset: Non-normalized (counts) distribution of the reversal times.

### 5. Cellular Potts model recapitulates quasi-periodic motion

The data presented so far are obtained from a large cell ensemble. Therefore, the distribution functions that describe the quasi-oscillatory cell motion present a robust and generic testbed for comparison with mathematical modeling. Recently, we have developed an extended Cellular Potts model that is capable of describing spatiotemporal dynamics of cells in 2D [30, 36]. In particular, we model the spatiotemporal dynamics of the contact area of a cell with a planar substrate, which is described by a set of discrete adhesion sites on a 2D lattice. With each configuration of the cell, which is characterized by a spreading area *A* and perimeter *P*, we associate the energy

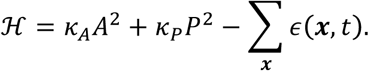

Here, the first two terms serve as a simplified description of the mechanical properties of cells [44-46]. In addition, the spatially resolved *polarization field, ϵ* (***x***, *t*) ∈[*ϵ*_0_ − Δ*ϵ*/2 … *ϵ*_0_ + Δ*ϵ*/2], emulates effective protrusive forces due to actin polymerization, actomyosin contractility and cell-substrate adhesions. The parameter *ϵ*_0_ denotes the average polarization field, and the parameter Δ*ϵ* denotes the polarization range. We assume that the cell gradually minimizes its configuration energy by adding or removing adhesion sites. In particular, one can illustrate that the cell is likely to protrude in regions with large polarization field and to retract in regions with small polarization field by interpreting *ϵ*(***x***, *t*) as an effective local *adhesion energy*. We simplify all intracellular signaling, which is mediated e.g. by the Rho GTPase family of proteins, into two prototypical feedback loops that break detailed balance. If the cell makes new contacts with the substrate (protrusion), then intracellular signaling reinforces the polarization field and therefore increases the likelihood of further protrusions. Then, the polarization field grows with a rate *μ* for all contact sites ***x*** that are surrounded by more protrusions than retractions within a fixed signaling range *R* [30, 36]:

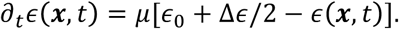

On the other hand, if the cell loses contacts with the substrate (retraction), then intracellular signaling weakens the polarization field and therefore increases the likelihood of further retractions. In particular, the polarization field decays with a rate *μ* for all contact sites ***x*** that are surrounded by more retractions than protrusions within a fixed signaling range *R* [30, 36]:

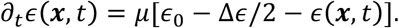

For all other contact sites between the cell and the substrate, the polarization field approaches a ‘rest state’ [30, 36]: *∂*_*t*_*ϵ*(***x***, *t*) = *μ*[*ϵ*_0_ – *ϵ*(***x***, *t*)]. This model reproduces persistent cell migration in a broad parameter regime [30, 36] and can therefore also reproduce oscillatory migration on microlanes. For a quantitative agreement between the migration timescales in our simulations and in our experiments, it is necessary to adjust the model parameters accordingly. However, it is a priori not clear whether there is a parameter set that can match the stochasticity of our experiments. Therefore, we use our experiments as a benchmark to test our model and to narrow down the parameters for the simulation of MDA-MB-231-cells. Finally, with these parameters, we investigate the lamellipodium morphology as it encounters different constrictions—a typical scenario for cell migration in vivo. In particular, we ask: how does the cell behave as it encounters differently shaped microlane tips?

To find a suitable parameter set, we adjusted the duration of a Monte Carlo step so that the average absolute velocity of migrating simulated cells, ⟨|*ν*|⟩ = 0.6 μ*m*/*min*, matches the experiments. Based on our previous work [30], we modified the parameters to account for cell persistence and stochasticity, and to achieve a sufficiently fine discretization of the cell body to resolve the micropattern tips. A detailed description of the parameters is provided in the Supporting Information S1. We then simulated individual cells with fixed parameters and constant average area on stripe-shaped microlanes. Note that this premise already marks a striking difference from the experiments: in the simulations we investigate a population of clonal and therefore identical cells, while the cells used in the experiments show a wide variation in morphology and migratory behavior. Therefore, we expect the simulations to underestimate all variances compared to the experiments.

We find that the model predicts large differences in cell shape and perimeter between polarized cells migrating in the microlanes and depolarized cells in the microlane tips, which qualitatively agrees with our experiments. However, cell size fluctuations in the model are very small, because we have kept the average polarization field *ϵ*_0_ constant. Interestingly, we find that the spreading area of the cells seems to shrink upon depolarization in the experiments. This might hint towards an additional regulation of cell-substrate adhesions that is currently not implemented in our model. Then, we investigated whether the model reproduces the correct statistics of quasi-oscillatory motion, and found both qualitative and quantitative agreement between the simulated cell trajectories and our experimental data (Fig 5). Here, we evaluated the distribution of cell reversal times in the simulations analogously to the experiments and observed a pronounced peak at 100 min (Fig 5a). Furthermore, the simulated cells show a similar spatial velocity distribution as in the experiments (Fig 5b). In addition, we also performed more intuitive comparisons between simulations and experiments. In particular, we find similar morphologies of (i) polarized and persistently migrating cells with a flat leading edge and a tapered rear, (ii) cells that depolarize after running into a dead end on their microlane, and (iii) repolarized cells (Fig 5c). To assess the actin distribution during cell migration, we transfected live cells with a fusion construct of the actin-binding peptide LifeAct and GFP (LifeAct-GFP). Intuitively comparing the representative kymographs for both a simulated cell and an experimental LifeAct-GFP transfected cell, both demonstrate similar oscillations on congruent microlanes (Fig 5d). In particular, we also find similarly sharp distributions of the polarization field in the model and the actin fluorescence intensity in our experiments (Fig 5d); in the model, this corresponds to a small signaling radius. However, we find that the cell also has a broad distribution of cortical actin in the cell body, which is not included in the model. The kymographs also show that there are several processes taking place during depolarization of the cells in experiments and in the simulation. The cytoskeletal activity at the leading edge is quenched relatively fast upon contact with the tip of the microlane, whereas the rear of the cell usually continues to move forward for some time. Taken together, our findings suggest that the quasi-periodic migration of cells on microlanes is well described by an extended Cellular Potts model. This model predicts cell polarization to robustly emerge from stochastic occurrence and subsequent self-reinforcement of cell protrusions, which then leads to a stochastic (re)polarization time. Therefore, the quasi-periodic oscillations observed in our experiments can emerge from a simple and generic coupling between cell polarization and cell movement.

**Fig 5.**
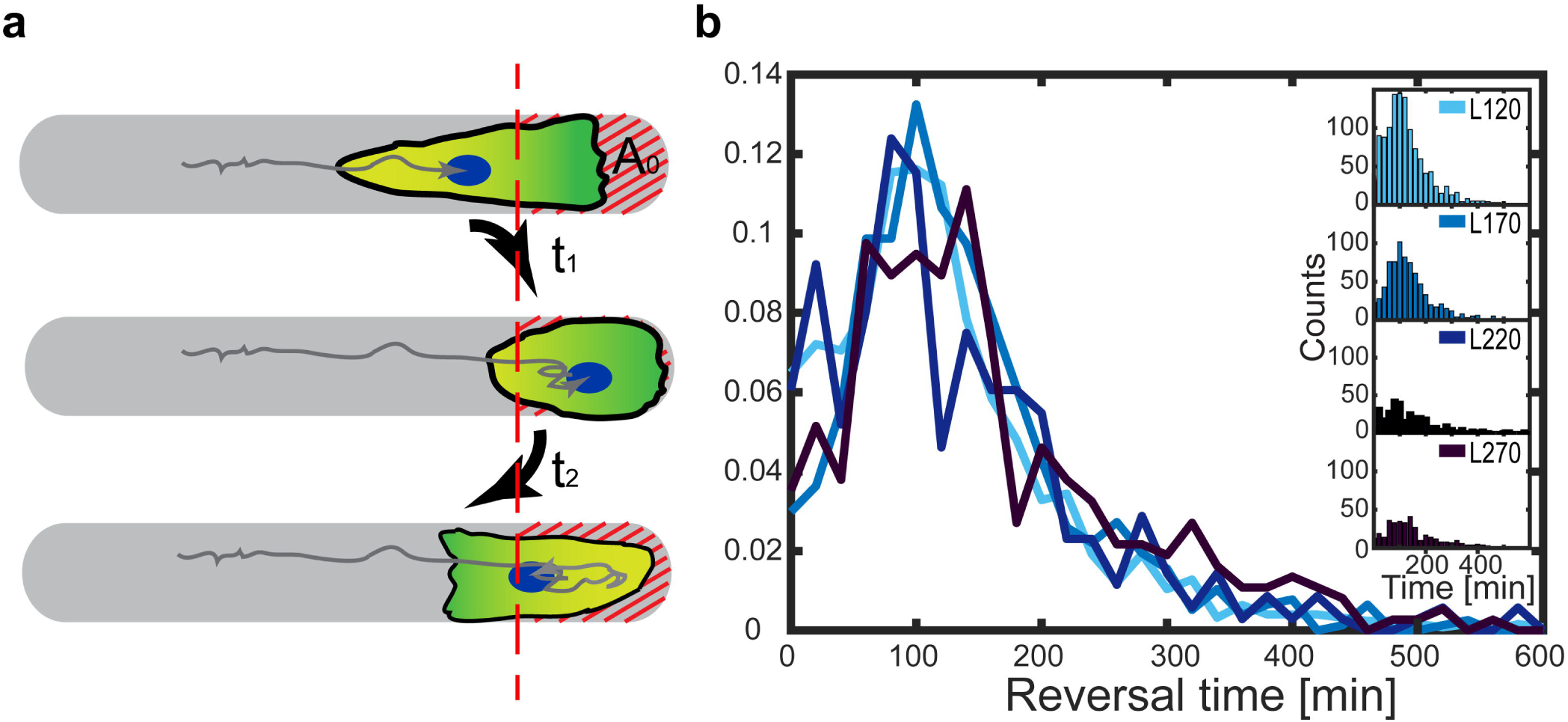
Comparison of computer simulations and experimental results. The microlane length is fixed at 170 μm with a round tip geometry. (a) Reversal time distributions of simulated and experimental cells. (b) Spatial mean absolute velocity distributions of simulated and experimental cells. (c) The extended Cellular Potts model features an internal polarization field. Our simulations reproduce the distinct run and rest phenotypes and yield cell center-of-mass trajectories that show quasi-periodic behavior (bottom); experimentally obtained cell trajectories are indicated on the upper right. (d) Comparison between the kymograph of a LifeAct-GFP transfected MDA-MB-231 cell with nuclear staining (bottom) and the kymograph of a simulated cell. Top: Zoom-in to a region that contains two periods of oscillation. Top left: Simulated cell. Top right: Experimental cell. Bottom: Zoom-out to the kymograph of an experimental cell that performs many periods of oscillation. Red arrows indicate the direction of cell motion. A video of the moving LifeAct-GFP transfected MDA-MB-231 cell can be found in Supporting Information “Video S4”.

### 6. Effect of curvature on cell depolarization

It is understood that the non-linear dynamics of actin polymerization and turnover depends on the cell shape and the geometry in which the cell migrates [23, 47]. In order to test the interplay between surface geometry and cell contour, we investigate the depolarization and repolarization of cells on microlanes with differently shaped tips. For example, a tapered tip allows us to explore how the reversal time depends on the deformation of the leading lamellipodium. To this end, we fabricated microlanes with four distinctly curved tips: round-, blunt-, sharp-, and concave-shaped, while keeping the total area and the width of the microlanes constant (Fig 6a) to assure the comparability of the cell behavior. Thus, depending on the tip geometry the length of the microlanes is slightly different to achieve a constant area. In order to observe many reversal events but also directed migration we established a length of 170 μm for the round tips. Exemplary fluorescent images of the microlanes are shown in Fig 6b. Across all studied geometries, we find that cells consistently exhibit oscillatory motion. Furthermore, the distribution of reversal times, or in other words, the depolarization-repolarization time, does not depend significantly on the respective tip geometry (Fig 6c). All reversal times are centered around approximately 100 min, which is consistent with the results in Fig 4b. A comparison between experiments and simulations, using the same algorithms for the analysis, yields similar reversal times (Fig 6d). However, the reversal time distribution is typically broader in the experiments than in the simulations, indicating that our simulations might underestimate the cell-to-cell variability.

**Fig 6.**
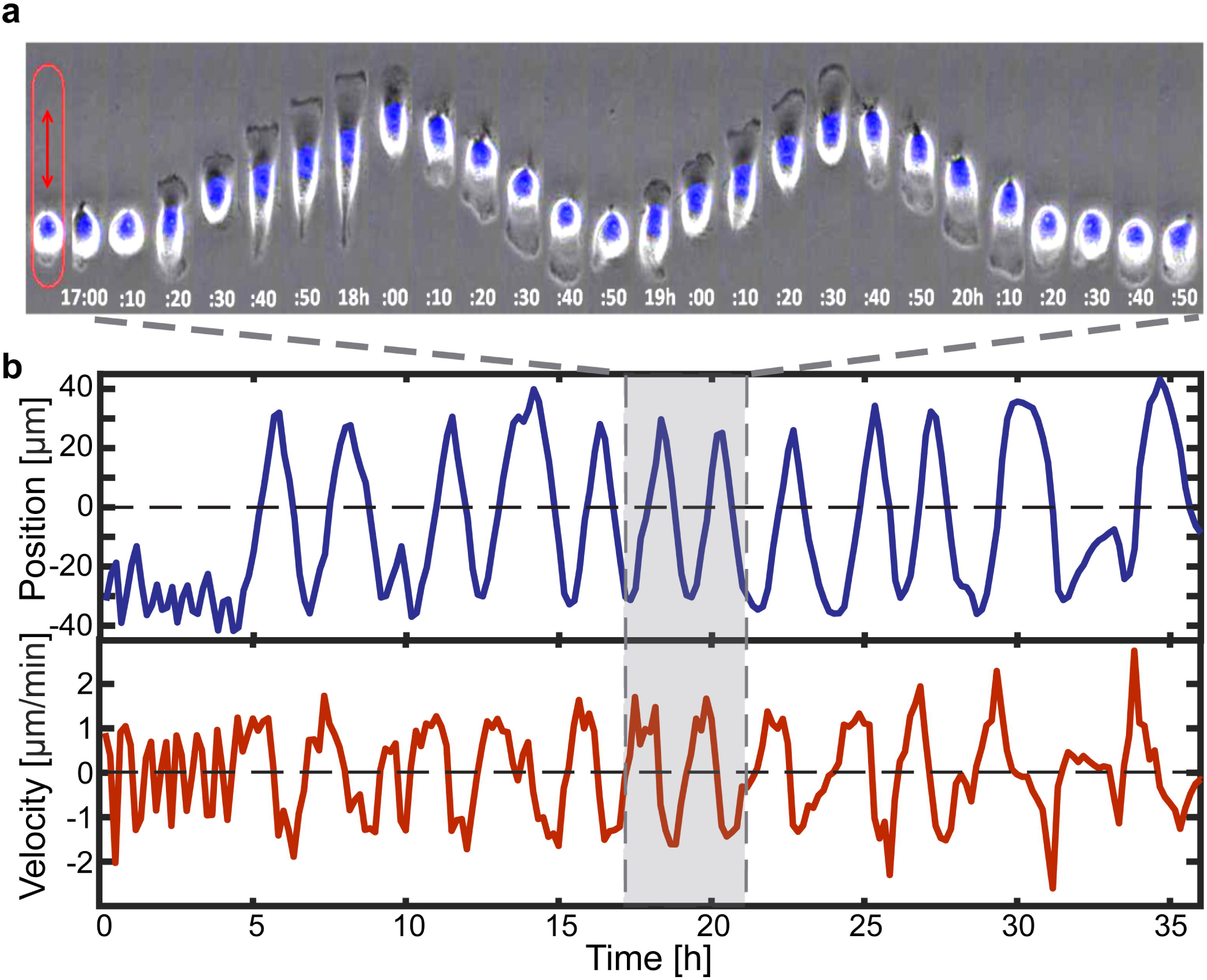
Effect of curvature on the repolarization time. (a) Schematic diagram of microlane geometry with four different tip shapes: blunt, concave, round, and sharp. The area of all microlanes and the reversal area A_0_ are kept constant. (b) Exemplary fluorescent images of microlanes of width W=20 μm. (c, d) Distribution of cell reversal times (Δt) when reaching the microlane tips, for four different tip shapes. (c) Experimental reversal time distribution. (d) Simulated reversal time distribution.

To gain additional insight into the spatiotemporal actin dynamics at the leading edge of the cell, we recorded a series of live time-lapse images of LifeAct-GFP transfected MDA-MB-231 cells. The deformation of the leading edge and the spatial distribution of F-actin in the protrusions are visualized in a close-up image series of the advancing lamellipodium at the microlane tips (Fig 7). There, we also show snapshots of the leading edge of simulated cells for a direct comparison between experiment and simulation. Both in experiment and simulation, we find that the lamellipodium splits in the concave-shaped ends showing local quenching of actin activity in the middle (Videos S5 and S6 in the Supporting Information). In contrast, in sharp ended microlanes the lamellipodium does not split but enters the sharp tip. Overall, the mesoscopic dynamics and reversal time of the cell do not seem to strongly depend on details of the microscopic dynamics of its lamellipodium. The constant repolarization time for all microlane lengths and tip shapes can be understood intuitively if the depolarization is fast compared to all other time scales and the repolarizing cell front does not have information about the tip geometry at the cell back. Furthermore, our experiments show that the PEGylated area does not constitute a solid mechanical boundary in our experiments. In many instances, we find transient actin protrusions into the PEGylated area. However, focal adhesion formation is impeded on the PEGylated substrate leading to a subsequent retraction. In contrast, in our computer simulations we strictly confine the cell contour to the micropattern. Furthermore, in the simulations, the contour of the advancing cell edge does not adopt the shape of the concave and sharp microlane tips to the same degree as in the experiments (Fig 7c and d). This might be due to (i) an overestimation of the perimeter stiffness or (ii) prohibiting the simulated cell from leaving the micropattern, which would allow less curved cell shapes.

**Fig 7.**
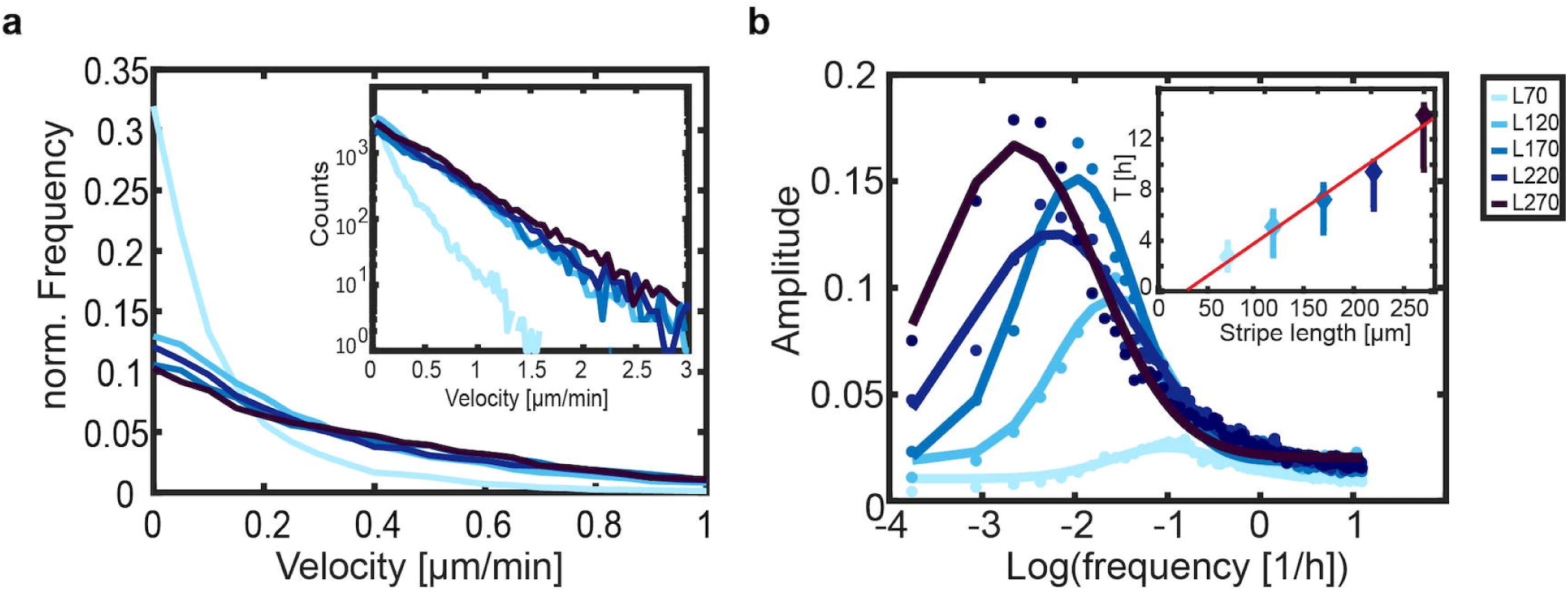
Geometry dependence of migratory arrest. Comparison between LifeAct GFP labeled MDA-MB-231 cells and simulated cells, which migrate towards differently shaped microlane tips (Lane length L = 170 μm, width W = 20 μm, 5 min time intervals). The top row (in green) shows fluorescence time-lapse data, while the bottom row shows the corresponding computer simulation for (a) round-shaped tips (b) blunt-shaped tips (c) sharp-shaped tips and (d) concave-shaped tips.

## Conclusion

In this work, we investigate single cells that migrate within short microlanes. In this form of confinement, cells exhibit a pole-to-pole migration mode. This behavior is quantified by the dominant oscillation frequency, the spatial distribution of cell positions and the persistent velocity of polarized migrating cells as a function of microlane length. The finding of quasi oscillatory pole-to-pole migration with repetitive depolarization-repolarization cycles is in agreement with previous measurements of the typical persistence length of directed migration on microtracks or microchannels, which was reported to be about 400 μm and hence larger than the length of the microlanes studied here [7, 32, 33, 41]. At the poles of the microlanes, actin polymerization in the leading protrusion is quenched which is most likely due to the reduced capability to form focal adhesions in the PEGylated area. In this way, further cell advancement is stopped. Subsequently, spontaneous protrusions form at the opposite, free edge of the cell and reverse the cell motion in the opposite direction. Interestingly the total reversal time does not depend on the length of the microlanes and hence appears to be independent of the migration history. In order to interrogate whether in-plane curvature of the tip boundary affects the spatio-temporal distribution of the actin polymerization front and possibly the reversal time, we constructed different tip shapes. We find that the tip shape has no significant influence on the macroscopic reversal times when the accessible area is preserved. However, we find that a concave tip shape leads to split protrusions showing local quenching of actin activity in the middle. In sharp microlanes, the lamellipodium does not split but enters the sharp tip. These experimental findings document that protrusions into constrictions are enhanced, while protrusions at concave interfaces are split. Our model provides a potential explanation of these experimental features. In the model, protrusions are amplified within a finite signaling radius, thereby coupling nearby protrusions. If the signaling radii are small compared to the microlane width (and have similar size as compared to the constrictions), then the cell can form two separate lamellipodia that invade the concave microlane tips. In between the split protrusions, the polarization field is quenched due to a lack of positive feedback.

We show that the experimentally observed pole-to-pole cell migration mode as well as the tip splitting at concave tips is recapitulated by an extended Cellular Potts model. Our model shows that the distribution of stochastic repolarization times might be explained as follows: stochastic membrane protrusions explore the vicinity of the cell. Then, if the cell can adhere in the explored region, these protrusions form stable lamellipodia through a self-reinforcing feedback loop. Note that our computational implementation conceptually resembles the protrusion fluctuation model used to describe directed cell motion on microratchets [38, 39]. Therefore, the general mechanism is the dynamics and reinforcement of exploring protrusions at both ends of the cell. For a further comparison between our extended Cellular Potts model and other models, including phase field approaches, we refer the reader to [30]. These phenomena emerge solely from the minimal feedback mechanism implemented in the Cellular Potts model; it does not currently take into account any structural details of the cytoskeleton. Further development of the Cellular Potts model to include myosin and the actin regulating Rho GTPases, which has been the subject of interest in analytical reaction-diffusion models of moving cells [7, 13, 15, 18], will possibly improve model predictions of the morphology and dynamics of the moving cell. In this context, the microlane assay proves useful as a testbed for future theory, facilitating the accumulation of statistics over repeated de- and repolarization events. As a cell depolarizes in different microlane geometries, the resulting spatial distribution of actin activity could lead to a better understanding of how cell adhesion and the local membrane curvature regulate actin polymerization [47-50]. In particular, computational models are challenged to recapitulate migratory behavior on various micro-pattern geometries in a consistent manner using an optimized, unique parameter set. Here, future studies combining cell migration assays on micropatterns and computational models will be valuable as a benchmark for model parameters. Hence, we propose to train physical models of cell migration on multiple experiments and in different confinements in order to gain predictive power. Such an approach will help to classify cytoskeleton dynamics and mechanisms that lead to distinct migration phenotypes.

## Materials and methods

### Micropatterning

Laser lithography. To prepare the master mold of the stamp for micropatterning, a silicon wafer was coated with TI Prime adhesion promoter and AZ40XT (MicroChemicals) photoresist. Areas for cell adhesion were exposed to UV light using laser direct imaging (Protolaser LDI, LPKF). The photoresist was developed (AZ 826 MIF, MicroChemicals) and then silanized (Trichloro(1H,1H,2H,2H-perfluorooctyl)silane, Sigma-Aldrich). To fabricate the stamp, polydimethylsiloxane (PDMS) monomer and crosslinker (DC 184 elastomer Kit, Dow Corning) were mixed in a 10:1 ratio (w/w), poured onto the master mold, and cured 3 h or overnight at 50 °C. The crosslinked PDMS layer was peeled off and manually cut into stamps.

Microcontact printing: Stripe-shaped microlanes were produced by microcontact printing. Firstly, PDMS stamps were exposed with UV-light (PSD-UV, Novascan Technologies) for 5 min. The stamps were then immersed in an aqueous solution of 40 μg/ml fibronectin (Yo Proteins) containing 10 μg/ml Alexa Fluor 488 dye (Life Technologies) labeled fibronectin for 45 min. The stamps were subsequently washed with ultrapure water. Stamps were dried under filtered airflow and then stamped onto a hydrophobic uncoated μ-Dish (Ibidi GmbH) bottom that underwent UV exposure for 15 min beforehand. The stamps were gently pressed with tweezers for a few seconds to ensure contact with the bottom. To then fabricate the cell-repelling areas, 30 μL of 2 mg/ml poly-L-lysine-grafted polyethylene glycol (PLL-g-PEG) (2 kDa PEG chains, SuSoS) dissolved in 10 mM Hepes and 150 mM NaCl solution was added. After the removal of the stamps, a glass cover slip was placed on the printed bottom to assure complete coverage with the PEG solution and then incubated for 30 min at room temperature. Finally, the printed bottom was washed with phosphate buffered saline (1x PBS) three times and stored in 1x PBS for further cell seeding. Unless otherwise specified, the patterns consisted of uniform stripes with a width of 20 μm.

### Cell culture and transfection

MDA-MB-231 breast cancer cells were cultured in modified Eagle’s medium (MEM-F10, c.c.pro) supplemented with 10% fetal calf serum (FCS, Invitrogen) and 2.5 mM L-glutamin (c.c.pro) at 37 °C in 5% CO2 atmosphere. For time-lapse phase-contrast images, cells were seeded at a density of 1 × 10^4^ cells per dish (μ-Dish, IBIDI). After 2 h, cell medium was replaced by 1 ml Leibovitz’s L-15 Medium (c.c.pro) containing 10% FCS and 25 nM Hoechst 33342 nucleic acid stain (Invitrogen) and incubated for 1 h at 37 °C before imaging.

For actin dynamics studies, seeded cells were further transfected with LifeAct-GFP mRNA. Briefly, ∼1× 10^4^ cells were seeded into a 35 mm μ-Dish and incubated 2 h at 37 °C in 5% CO2 for cell adhesion. 1.25 μl Lipofectamine MessengerMax Reagent (Invitrogen) was diluted in 123.75 μl OptiMEM (Life Technologies) transfection medium and incubated 10 min at room temperature. 500 ng mRNA (0.5 μl × 1000 ng/μl) was diluted in 124.5 μl OptiMEM. Both solutions were mixed and incubated for 5 min at room temperature for lipoplex formation. Adhered cells were washed with 1x PBS, and carefully added to the 250 μl transfection mix. After a 1 h incubation at 37 °C in 5% CO2, the cell transfection mix was replaced by 1 ml Leibovitz’s L-15 Medium (c.c.pro) containing 10% FCS before proceeding to time lapse imaging.

Laboratory protocols can be found: http://dx.doi.org/10.17504/protocols.io.bcdiis4e

### Live cell imaging and microscopy

For migration studies, scanning time lapse measurements were acquired using an automated inverted microscope iMIC (Till Photonics). The microscope was equipped with a 10x Zeiss objective and a 40x Zeiss objective, an ORCA-03G camera (HAMAMATSU), and an Oligochrome lamp (Till Photonics). During the measurements, cells were maintained in L-15 Meidium containing 10% FCS at 37 °C using a temperature-controlled mounting frame (Ibidi temperature controller, Ibidi). Phase contrast and fluorescent images were automatically acquired at 10 min intervals, unless noted otherwise. To analyze the actin dynamics at the interface of micropatterns, images were acquired at intervals between 20 s - 1 min.

### Image processing and data analysis

Image analysis was carried out using ImageJ (National Institutes of Health, NIH). Images of isolated cells migrating in the stripe-shaped microlanes were first manually cropped. The trajectory of each stained nucleus was preprocessed by first applying a bandpass filter and a threshold to the fluorescence images, and the geometric center of mass of the nucleus was subsequently evaluated. The geometric mean of the nucleus position was used as a proxy for the cell position. Only single-cells that explored the whole stripe where analyzed. Cell tracks were excluded from further analysis in the following cases: cell tracks shorter than 36 h due to cell division or spanning out of the micropattern, and tracks of non-moving or dead cells (less than 5%).

Trajectories of individual cells were analyzed in Matlab. Only the component of the cell position in the direction of the corresponding microlane was considered resulting in a 1D trajectory. The center of the microlane was determined by taking the average of the two points where cells got closest to each tip of the microlane.

## Supporting information

**Table S1. Parameters used for the computational simulation in this work**.

**Table S2. Number of analyzed cells for different lengths of microlanes with round tips**.

**Table S3. Number of analyzed cells for microlanes with different geometric tips**.

**Fig S1. Mean cell velocity as a function of the distance to the nearest tip in microlanes of different lengths**.

**Fig S2. Additional exemplary trajectories of cells with different frequencies on the microlane L=120 μm**.

**Video S1. A cell (MDA-MB-231) migrates on a stripe-shaped micropattern (L = 120 μm)**.

**Video S2. An exemplary cell migrates on a shorter microlane (L = 70 μm)**.

**Video S3. An exemplary cell migrates on a longer microlane (L = 170 μm)**.

**Video S4. A LifeAct-GFP transfected cell migrates on the microlane L = 170 μm**.

**Video S5. Actin dynamics of a LifeAct-GFP transfected cell on a microlane with concave-shaped tips**.

**Video S6. Actin dynamics of a LifeAct-GFP transfected cell on a microlane with sharp-shaped tips**.

**Data S1. Data of all cells used for this work in csv format. Time given in minutes and “koor” specifies the distance from the center of the stripe in micrometers along the long axis of the pattern, i**.**e. the direction of migration**.

## References

1. Mitchison TJ, Cramer LP. Actin-Based Cell Motility and Cell Locomotion. Cell. 1996;84(3):371–9.

2. Keren K, Pincus Z, Allen GM, Barnhart EL, Marriott G, Mogilner A, et al. Mechanism of shape determination in motile cells. Nature. 2008;453(7194):475–80.

3. Kruse K, Camalet S, Julicher F. Self-propagating patterns in active filament bundles. Phys Rev Lett. 2001;87(13). PubMed PMID: WOS:000171219500059.

4. Carlsson AE. Dendritic Actin Filament Nucleation Causes Traveling Waves and Patches. Phys Rev Lett. 2010;104(22):228102.

5. Giannone G, Dubin-Thaler BJ, Dobereiner HG, Kieffer N, Bresnick AR, Sheetz MP. Periodic lamellipodial contractions correlate with rearward actin waves. Cell. 2004;116(3):431-43. PubMed PMID: WOS:000188825800011.

6. Bretschneider T, Anderson K, Ecke M, Müller-Taubenberger A, Schroth-Diez B, Ishikawa-Ankerhold HC, et al. The Three-Dimensional Dynamics of Actin Waves, a Model of Cytoskeletal Self-Organization. Biophysical Journal. 2009;96(7):2888–900.

7. Maiuri P, Rupprecht J-F, Wieser S, Ruprecht V, Bénichou O, Carpi N, et al. Actin Flows Mediate a Universal Coupling between Cell Speed and Cell Persistence. Cell. 2015;161(2):374–86.

8. Shao D, Levine H, Rappel W-J. Coupling actin flow, adhesion, and morphology in a computational cell motility model. Proceedings of the National Academy of Sciences. 2012;109(18):6851–6. doi: 10.1073/pnas.1203252109.

9. Allard J, Mogilner A. Traveling waves in actin dynamics and cell motility. Current Opinion in Cell Biology. 2013;25(1):107–15.

10. Verkhovsky AB. The mechanisms of spatial and temporal patterning of cell-edge dynamics. Curr Opin Cell Biol. 2015;36 :113–21. doi: 10.1016/j.ceb.2015.09.001. PubMed PMID: 26432504.

11. Lawson CD, Ridley AJ. Rho GTPase signaling complexes in cell migration and invasion. J Cell Biol. 2018;217(2):447–57.

12. Goryachev AB, Pokhilko AV. Computational model explains high activity and rapid cycling of Rho GTPases within protein complexes. PLoS computational biology. 2006;2(12):e172.

13. Mori Y, Jilkine A, Edelstein-Keshet L. Wave-pinning and cell polarity from a bistable reaction-diffusion system. Biophysical journal. 2008;94(9):3684–97.

14. Klünder B, Freisinger T, Wedlich-Söldner R, Frey E. GDI-mediated cell polarization in yeast provides precise spatial and temporal control of Cdc42 signaling. PLoS computational biology. 2013;9(12):e1003396.

15. Edelstein-Keshet L, Holmes WR, Zajac M, Dutot M. From simple to detailed models for cell polarization. Philosophical Transactions of the Royal Society B: Biological Sciences. 2013;368(1629):20130003.

16. Bement WM, Leda M, Moe AM, Kita AM, Larson ME, Golding AE, et al. Activator–inhibitor coupling between Rho signalling and actin assembly makes the cell cortex an excitable medium. Nature cell biology. 2015;17(11):1471.

17. Falcke M. Concentration profiles of actin-binding molecules in lamellipodia. Physica D: Nonlinear Phenomena. 2016;18-319:50–7.

18. Byrne Kate M, Monsefi N, Dawson John C, Degasperi A, Bukowski-Wills J-C, Volinsky N, et al. Bistability in the Rac1, PAK, and RhoA Signaling Network Drives Actin Cytoskeleton Dynamics and Cell Motility Switches. Cell Systems. 2016;2(1):38–48.

19. Halatek J, Brauns F, Frey E. Self-organization principles of intracellular pattern formation. Philosophical Transactions of the Royal Society B: Biological Sciences. 2018;373(1747):20170107.

20. Brauns F, Halatek J, Frey E. Phase-space geometry of reaction--diffusion dynamics. arXiv preprint 181208684. 2018.

21. Halatek J, Frey E. Rethinking pattern formation in reaction–diffusion systems. Nature Physics. 2018;14(5):507.

22. Marée AF, Jilkine A, Dawes A, Grieneisen VA, Edelstein-Keshet L. Polarization and movement of keratocytes: a multiscale modelling approach. Bulletin of mathematical biology. 2006;68(5):1169–211.

23. Marée AFM, Grieneisen VA, Edelstein-Keshet L. How Cells Integrate Complex Stimuli: The Effect of Feedback from Phosphoinositides and Cell Shape on Cell Polarization and Motility. PLOS Computational Biology. 2012;8(3):e1002402. doi: 10.1371/journal.pcbi.1002402.

24. Mogilner A, Keren K. The Shape of Motile Cells. Curr Biol. 2009;19(17):R762–R71.

25. Ziebert F, Aranson IS. Effects of adhesion dynamics and substrate compliance on the shape and motility of crawling cells. PloS one. 2013;8(5):e64511.

26. Ziebert F, Swaminathan S, Aranson IS. Model for self-polarization and motility of keratocyte fragments. Journal of The Royal Society Interface. 2011;9(70):1084–92.

27. Camley BA, Zhao Y, Li B, Levine H, Rappel W-J. Periodic Migration in a Physical Model of Cells on Micropatterns. Phys Rev Lett. 2013;111(15). doi: 10.1103/PhysRevLett.111.158102.

28. Albert PJ, Schwarz US. Dynamics of cell shape and forces on micropatterned substrates predicted by a cellular Potts model. Biophysical journal. 2014;106(11):2340–52.

29. Goychuk A, Brückner DB, Holle AW, Spatz JP, Broedersz CP, Frey E. Morphology and Motility of Cells on Soft Substrates. arXiv preprint 180800314. 2018.

30. Thueroff F, Goychuk A, Reiter M, Frey E. Bridging the gap between single cell migration and collective dynamics. doi: 10.7554/eLife.46842

31. Lautenschläger F, Piel M. Microfabricated devices for cell biology: all for one and one for all. Current opinion in cell biology. 2013;25(1):116–24.

32. Maiuri P, Terriac E, Paul-Gilloteaux P, Vignaud T, McNally K, Onuffer J, et al. The first World Cell Race. Current Biology. 2012;22(17):R673–R5.

33. Wilson K, Lewalle A, Fritzsche M, Thorogate R, Duke T, Charras G. Mechanisms of leading edge protrusion in interstitial migration. Nature Communications. 2013;4 :2896. doi: 10.1038/ncomms3896

34. Doyle AD, Wang FW, Matsumoto K, Yamada KM. One-dimensional topography underlies three-dimensional fibrillar cell migration. The Journal of cell biology. 2009;184(4):481–90.

35. Picone R, Ren X, Ivanovitch KD, Clarke JDW, McKendry RA, Baum B. A Polarised Population of Dynamic Microtubules Mediates Homeostatic Length Control in Animal Cells. PLOS Biology. 2010;8(11):e1000542. doi: 10.1371/journal.pbio.1000542.

36. Segerer FJ, Thüroff F, Alberola AP, Frey E, Rädler JO. Emergence and persistence of collective cell migration on small circular micropatterns. Phys Rev Lett. 2015;114(22):228102.

37. Mahmud G, Campbell CJ, Bishop KJM, Komarova YA, Chaga O, Soh S, et al. Directing cell motions on micropatterned ratchets. Nature Physics. 2009;5 :606. doi: 10.1038/nphys1306

38. Caballero D, Voituriez R, Riveline D. The cell ratchet: Interplay between efficient protrusions and adhesion determines cell motion. Cell Adhesion & Migration. 2015;9(5):327–34. doi: 10.1080/19336918.2015.1061865.

39. Caballero D, Comelles J, Piel M, Voituriez R, Riveline D. Ratchetaxis: Long-Range Directed Cell Migration by Local Cues. Trends in Cell Biology. 2015;25(12):815–27.

40. Brückner DB, Fink A, Schreiber C, Röttgermann PJ, Rädler JO, Broedersz CP. Stochastic nonlinear dynamics of confined cell migration in two-state systems. Nature Physics. 2019;15(6):595.

41. Schreiber C, Segerer FJ, Wagner E, Roidl A, Rädler JO. Ring-Shaped Microlanes and Chemical Barriers as a Platform for Probing Single-Cell Migration. Scientific Reports. 2016;6 :26858. doi: 10.1038/srep26858

42. Zhang J, Guo WH, Wang YL. Microtubules stabilize cell polarity by localizing rear signals. Proceedings of the National Academy of Sciences of the United States of America. 2014;111(46):16383–8. doi: 10.1073/pnas.1410533111. PubMed PMID: 25368191; PubMed Central PMCID: PMC4246331.

43. Fraley SI, Feng Y, Giri A, Longmore GD, Wirtz D. Dimensional and temporal controls of three-dimensional cell migration by zyxin and binding partners. Nat Commun. 2012;3 :719. doi: 10.1038/ncomms1711. PubMed PMID: 22395610.

44. Daub JT, Merks RMH. A Cell-Based Model of Extracellular-Matrix-Guided Endothelial Cell Migration During Angiogenesis. Bulletin of Mathematical Biology. 2013;75(8):1377–99. doi: 10.1007/s11538-013-9826-5.

45. Kabla AJ. Collective cell migration: leadership, invasion and segregation. Journal of The Royal Society Interface. 2012;9(77):3268–78. doi: doi:10.1098/rsif.2012.0448.

46. Merks RMH, Glazier JA. A cell-centered approach to developmental biology. Physica A: Statistical Mechanics and its Applications. 2005;352(1):113–30.

47. James J, Goluch ED, Hu H, Liu C, Mrksich M. Subcellular curvature at the perimeter of micropatterned cells influences lamellipodial distribution and cell polarity. Cell motility and the cytoskeleton. 2008;65(11):841–52.

48. Jeon H, Koo S, Reese WM, Loskill P, Grigoropoulos CP, Healy KE. Directing cell migration and organization via nanocrater-patterned cell-repellent interfaces. Nature materials. 2015;14(9):918.

49. Jiang X, Bruzewicz DA, Wong AP, Piel M, Whitesides GM. Directing cell migration with asymmetric micropatterns. Proceedings of the National Academy of Sciences. 2005;102(4):975–8.

50. Hsu H-F, Bodenschatz E, Westendorf C, Gholami A, Pumir A, Tarantola M, et al. Variability and Order in Cytoskeletal Dynamics of Motile Amoeboid Cells. Phys Rev Lett. 2017;119(14):148101. doi: 10.1103/PhysRevLett.119.148101.

